# The SWI/SNF and RSC Cooperatively Remodel the Promoters of Unfolded Protein Response Targets and Heat Shock Genes

**DOI:** 10.1101/2021.05.09.443345

**Authors:** Rakesh Kumar Sahu, Sakshi Singh, Raghuvir Singh Tomar

**Affiliations:** Laboratory of Chromatin Biology, Department of Biological Sciences, Indian Institute of Science Education and Research Bhopal, India

**Keywords:** SWI/SNF, RSC, Chromatin remodelling complexes, ER stress, UPR, Transcription regulation

## Abstract

The ATP-dependent chromatin remodelling complexes maintain the chromatin dynamics, enabling the gene expression or its silencing. The SWI/SNF subfamily remodelers (SWI/SNF and RSC) generally promote gene expression by displacing or evicting nucleosomes at the promoter regions. Their action creates a nucleosome-depleted region where transcription machinery accesses the DNA. Their involvement has been shown critical for the induction of stress-responsive transcription programs. Although the role of SWI/SNF and RSC complexes in transcription regulation of heat shock responsive genes is well studied, their involvement at other pathway genes such as UPR, HSP and PQC is less known. In this study, we showed that the SWI/SNF occupies promoters of UPR, HSP and PQC genes in response to the unfolded protein stress, and its recruitment at UPR promoters is dependent on the Hac1 transcription factor and other epigenetic factors like Ada2 and Ume6. Disruption of SWI/SNF’s activity does not affect the remodelling of these promoters or gene expression. However, inactivation of both RSC and SWI/SNF complexes diminishes expression of most of the UPR, HSP and PQC genes tested. Altogether these results suggest that these two remodelers work together or one compensates the loss of the other to ensure optimal induction of the stress-responsive genes.

## Introduction

The cellular proteome is continuously threatened by environmental and physiological stress like heat, exposure to toxic chemicals and heavy metals (such as cadmium and arsenic), nutrient starvation and senescence (1–7). Under such stresses, proteostasis mechanisms, like, protein folding, protein trafficking, and the clearance of undesirable or terminally unfolded proteins get compromised (3, 5, 8, 9). This ultimately leads to the accumulation of misfolded proteins in the cytoplasm, and in response, cells activate multiple signalling pathways, generally intended to activate the stress-responsive transcriptional programs and alleviate the stress (3, 10–12). The unfolded protein response (UPR) is triggered when the accumulated unfolded or improperly folded proteins in the ER lumen are sensed by the ER transmembrane endonuclease protein-Ire1 (13, 14). Activated Ire1 transduce the UPR signalling by performing non-conventional splicing of *HAC1* mRNA encoding for a basic leucine zipper (bZIP) transcription factor, relieving its translational attenuation (15–17). Upon translation, the Hac1 transcription factor is imported to the nucleus, where it activates its target genes by recognizing an unfolded protein response element (UPRE) sequence present on the promoters (15, 18). Its targets include genes involved in protein folding, protein modification (glycosylation), ER-associated degradation (ERAD), ER-to-Golgi transport and lipid metabolism (19, 20).

Several UPR-independent transcription factors such as heat shock factor (HSF), Msn2/4 and Rpn4 also respond to, and relieve, unfolded protein stress (21–26). In the absence of stress, HSF1 remains bound with Hsp90, Hsp70, and the Hsp70 co-chaperone- Hsp40, and prevented from forming its trimeric form (27–32). Upon heat stress, HSF1 trimerizes and activates chaperone genes by binding on heat-shock elements (HSEs) present on their promoter regions (23, 33–35). Similarly, Msn2/4 factors bind to stress response element (STRE) sequences on target genes (36) and promote their transcription initiation in response to multiple stress conditions, including nutrient starvation, heat shock, lipotoxicity, osmotic- and oxidative stresses (12, 21, 22, 37–41). Besides HSF1 and Msn2/4, Rpn4 activates genes encoding components of the ubiquitin-proteasome pathway to maintain protein quality in response to ER stress, which further complements UPR signalling (26).

Chromatin remodelling of the gene promoters is one of the initial and important events for stress-inducible gene transcription (42–45). In general, mobilization of nucleosomes, or eviction of histone octamers near the transcription start site (TSS), and upstream of it, allows transcription machinery to access and bind to the target gene regulatory sites (such as upstream activation sequence (UAS) and TATA Box) and initiate its transcription (reviewed in (46)). While the HSF1 actively participates in chromatin remodelling at its target gene promoters (47), the involvement of other transcription factors, Hac1, Rpn4 and Msn2/4, in this process is not well understood. Histone modifications, especially H3 and H4 acetylation, accompany the remodelling event at HSF1 targets and are believed to help recruit SWI/SNF chromatin remodelling complex on to the targets (48). This megadalton complex alters chromatin by mobilizing nucleosomes using its ATPase–translocase ‘motor’ (49–51). Unlike ISWI and CHD remodelers, which restores a closed chromatin state by arranging nucleosomes in a regularly spaced array (52–58), SWI/SNF primarily functions to open the chromatin region by displacing or evicting nucleosomes so that transcription machinery and DNA repair factors can bind to the DNA (50, 59, 60). Mechanistically, the catalytic subunit of SWI/SNF- Snf2 translocates the nucleosomal-DNA by continuously breaking the histone-DNA interactions with the help of two RecA like lobes in its ATPase domain. Continued DNA translocation over a particular histone octamer creates a sliding effect on the nucleosome and increase the distance between nucleosomes. Furthermore, with continued DNA translocation, SWI/SNF can spool off DNA from the neighbouring nucleosome, and it can dislodge the DNA from the substrate nucleosome with a high translocation efficiency; either way, it forms an open chromatin region (reviewed in (61, 62)). This nucleosome-depleted region (NDRs) created by SWI/SNF at the TSS of a gene often positively impacts transcription initiation (56, 63). Deletion of *SNF2* delays histone eviction from some HSF1 target genes during heat-induced expression, affecting HSF1 binding and RNA polymerase II recruitment (64, 65). Additionally, it eliminates the transcription induced degradation of Msn2 at the *HSP12* promoter during heat stress. Altogether, these observations indicate that HSF1 and Msn2/4 require a prior chromatin remodelling by SWI/SNF to act on their target genes (64).

SWI/SNF is known to cooperate with multiple cofactors to activate various stress-responsive transcriptional programs. For instance, in amino acid deprivation condition, SWI/SNF acts on a subset of genes strongly induced by Gcn4, in cooperation with Gcn5 histone acetyltransferase (HAT) and the Hsp70 co-chaperone-Ydj1 (66). SWI/SNF also functions in concert with RSC, a member of the SWI/SNF family of chromatin remodelling complexes, in repositioning the -1 and +1 nucleosomes of Gcn4-induced genes in amino acid deprivation (63). Here SWI/SNF function more prominently than RSC on some highly remodelled promoters in sliding -1 and +1 nucleosomes, while RSC acts on a majority of Gcn4 induced genes to maintain proper NDR widths (63). Similar functional cooperation among SWI/SNF, RSC and other cofactors was also observed on a subset of constitutively expressed genes, implying a critical role of these remodelling enzymes at highly expressed genes (63, 66). A key question about SWI/SNF is how its recruitment at stress-inducible promoters is regulated. Previous studies have explored this process which shows that the Sko1-Cyc8-Tup1 repressor complex and Rlm1 transcription factor recruits SWI/SNF on some stress-responsive genes in response to osmotic- and cell wall stresses, respectively (67, 68). Further, SWI/SNF recruitment over a DNA damage-inducible gene-*RNR3* requires general transcription factor TFIID and RNA polymerase II in genotoxic stress (69).

Although the remodelling activity of SWI/SNF at the heat shock and stress-responsive genes is extensively studied (48, 63–66, 68, 70, 71), its role in the induction of UPR genes is unknown. Further, multiple chaperones and protein quality control genes get activated in response to the unfolded protein stress, and the role of SWI/SNF in activating these genes is less understood. In this study, we examined the involvement of SWI/SNF in the induction of UPR, HSP and PQC genes in tunicamycin and cadmium-induced ER stress. Our results suggest SWI/SNF occupies UPR, HSP and PQC gene promoters in ER stress and Hac1, Ada2 and Ume6 facilitate its recruitment at UPR gene promoters. However, the disruption of SWI/SNF activity does not affect the expression of these genes. Furthermore, disruption of both SWI/SNF and RSC activity affected induction of a majority of genes tested, suggesting a functional cooperation between these two remodelling complexes.

## Results

### SWI/SNF is recruited to UPR gene promoters in ER stress but dispensable for gene expression

To examine the SWI/SNF complex’s involvement in remodelling the ER-stress induced genes, we first measured the SWI/SNF occupancy over the promoters in cadmium and tunicamycin treatments. The genes analyzed include UPR regulated genes (*EUG1*, *KAR2*, *ERO1*, *PDI1*, *SCJ1*, *FKB2* and *MCD4*); heat shock genes (*SSA4*, *HSP12*, *HSP42* and *HSP82*); genes involved in protein quality control (*RPN4*, *UBI4* and *CDC48*); and other stress-responsive genes (*CTT1* and *RAD59*). We monitored the SWI/SNF enrichment at activator binding sites (UPRE and TATA box) of these genes to know the involvement of SWI/SNF in preinitiation complex formation.

Our Snf2-ChIP results show a substantial increase of SWI/SNF binding over *KAR2* in tunicamycin induced ER stress (Figure 1A). While, the fold enrichment of SWI/SNF over the *PDI1*, *EUG1*, *ERO1*, *MCD4* and *FKB2* promoters is around 1.8-2- fold in tunicamycin-induced condition compared with the unstressed condition (Figure 1A). The *SCJ1* promoter showed a very slight increment of SWI/SNF binding in tunicamycin stress (Figure 1A).

**Figure 1.**
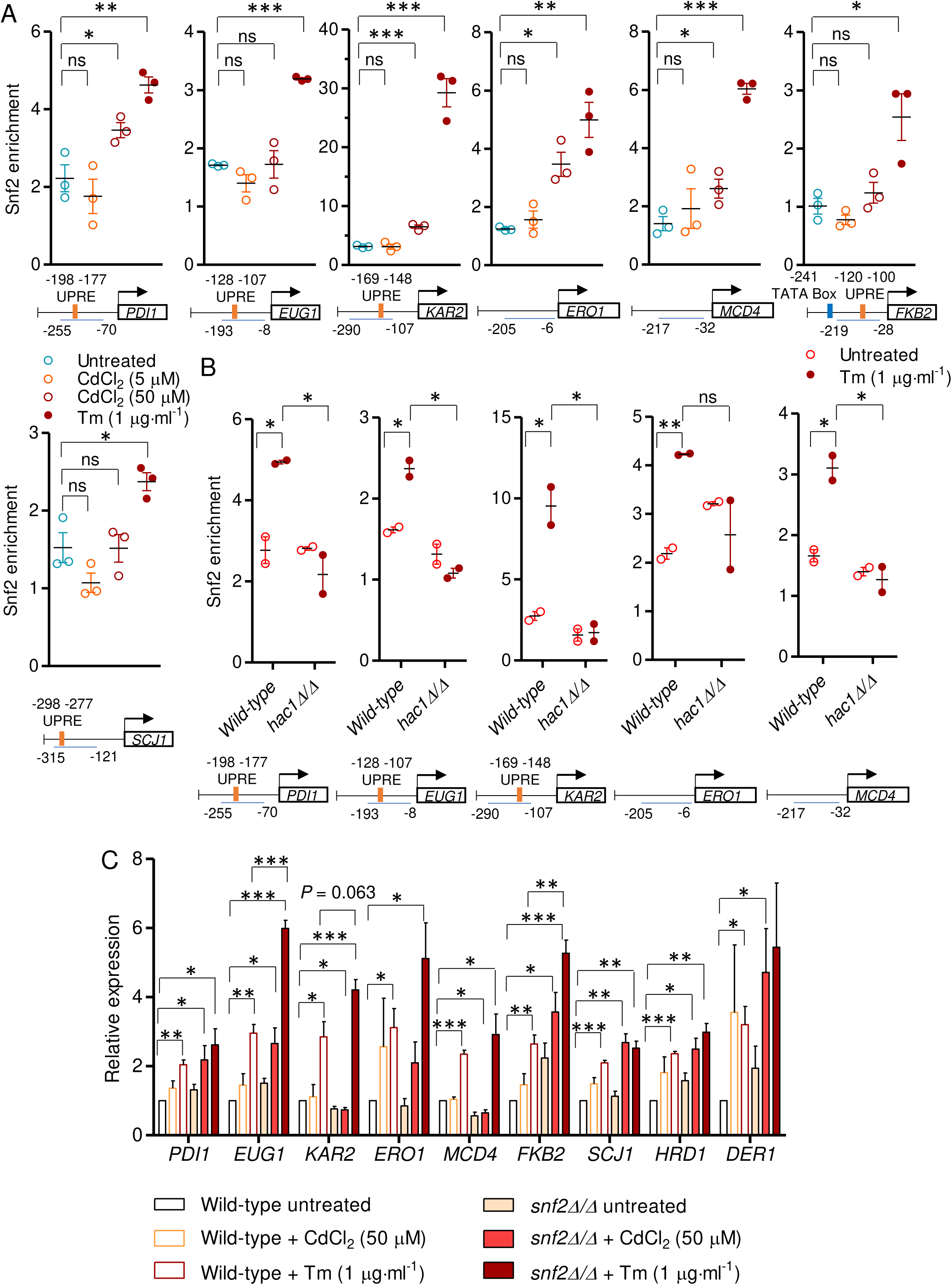
SWI/SNF complex is recruited over UPR gene promoters in ER stress yet dispensable for gene expression. (A) The fold enrichment of Snf2 over the UPR gene promoters are shown. The location of the TATA box, UPRE and the amplicon size of the promoters are indicated below each panel. Values shown are the Snf2 fold enrichment and the mean of three independent biological repeats (n = 3). Error bars indicate the standard error of the mean (SEM). Student’s *t*-test statistical analyses were performed, and significant differences are indicated as; *, *P* ≤ 0.05; **, *P* ≤ 0.005; ***, *P* ≤ 0.001; ns, non-significant. (B) The Snf2 fold enrichment over UPR target genes in wild-type and *hac1Δ/Δ* cells are shown. The values shown are the fold enrichment and the mean ± SEM of two independent biological repeats (n = 2). Student’s *t*-test statistical analyses were performed, and significant differences are indicated as; *, *P* ≤ 0.05; **, *P* ≤ 0.005; ns, non-significant. (C) The expression of UPR target genes in wild-type and *snf2Δ/Δ* cells under ER stress are shown relative to the transcript level of unstressed wild-type cells. Values shown are the mean and SEM of three independent biological repeats (n = 3). Student’s *t*-test statistical analyses were performed, and significant differences are indicated as; *, *P* ≤ 0.05; **, *P* ≤ 0.005; ***, *P* ≤ 0.001; ns, non-significant.

The *KAR2*, *PDI1* and *ERO1* are among the highly induced genes in ER-stress, which code for factors involved in protein folding in the ER lumen. The Pdi1 and Ero1 catalyze disulfide bond formation in newly folding proteins using the cofactor flavin adenine dinucleotide (FAD), while Kar2 binds to newly forming peptides inside the ER lumen and facilitate their translocation into the ER and prevent their misfolding. The rest of the UPR regulated genes tested encode protein disulfide isomerase (Eug1), peptidyl-prolyl cis-trans isomerase (Fkb2), DnaJ like chaperone (Scj1) and protein involved in glycosylphosphatidylinositol (GPI) anchor synthesis (Mcd4), all involved in protein modification and folding. The SWI/SNF occupies the UPRE at the UPR gene promoters during induction, which indicates that SWI/SNF mediated remodelling of the UPRE region might be required for Hac1 action at these promoters.

In cadmium-induced ER stress, the enrichment of SWI/SNF is slightly increased over *PDI1*, *KAR2*, *ERO1*, *MCD4* and *FKB2,* and its enrichment at these promoters is almost negligible in the acute cadmium stress compared to the unstressed condition (Figure 1A). Altogether, these results suggest that SWI/SNF binds and remodels the UPR induced genes in response to ER stress. The recruitment of SWI/SNF at the UPR promoters is Hac1 driven, as the deletion of *HAC1* reduces the SWI/SNF occupancy at these promoters (Figure 1B). This suggests that the Hac1 transcription factor cooperates with SWI/SNF or other chromatin remodelling factors to activate UPR targets.

To assess the necessity of the SWI/SNF complex in ER stress-induced UPR gene induction, we monitored the UPR gene induction in the *SNF2* deletion condition. In the wild-type cells, although there was a slight or no increase in UPR gene expression in cadmium stress relative to the unstressed condition, there was a substantial increase of UPR gene induction (2-4 folds) in tunicamycin induced ER stress condition (Figure 1C), correlating with the Snf2 enrichment at these promoters. However, to our surprise, the expression of UPR genes in ER stress was not affected in *SNF2* deleted strain (Figure 1C). Indeed, the expression of some genes like *EUG1*, *KAR2*, *ERO1* and *FKB2* in tunicamycin stress is higher in *snf2Δ/Δ* cells compared to that of wild-type cells (Figure 1C). Thus, the gene expression analysis indicates that, although SWI/SNF is recruited at UPR promoters in response to the ER stress, its absence can be compensated by other chromatin remodelling factors making the SWI/SNF dispensable for gene expression.

### SWI/SNF binding over UPR gene promoters depends on the Ada2 and Ume6

The post-translational modifications at the histones regulate the stress-induced transcriptional program (48, 72–74). Modified histone residues relay the gene activatory or repressive signals by acting as a docking site or recognition signal for many transcription activators and chromatin-modifying enzymes, including SWI/SNF (75). To understand the role of histone modification in SWI/SNF’s recruitment over UPR-responsive and UPR-independent promoters in ER stress, we looked at the SWI/SNF abundance in the absence of some histone-modifying enzymes-Ada2, Ume6 and Set2. The Ada2, with two other subunits Ada3 and Gcn5, forms the catalytic core of ADA and SAGA HAT complexes, where it potentiates the acetylase activity of Gcn5 (76). The Ume6, on the contrary, is a component of the Rpd3L histone deacetylase (HDAC) complex, generally involved in transcriptional repression of genes associated with meiosis, phospholipid biosynthesis, nitrogen metabolism and heat shock (77–83). Besides histone deacetylation, Ume6 physically interacts with Isw2 chromatin remodelling complex and targets Isw2 genome-wide by DNA looping mechanism and orchestrates the transcriptional repression (84, 85). We found that disruption of Ada2 and Ume6 histone-modifying components affect SWI/SNF recruitment at ER stress responsive promoters.

The binding of Snf2 over *PDI1*, *EUG1* and *MCD4* is reduced in the absence of Ada2 and Ume6, suggesting their activity is required for binding of SWI/SNF over these promoters (Figure 2A). Further, a drastic reduction of Snf2 binding was seen over *KAR2* promoter in *ada2Δ/Δ* and *ume6Δ/Δ* mutants, indicating that the action of Ada2 and Ume6 is necessary for the recruitment of SWI/SNF over *KAR2* (Figure 2A). Additionally, *ADA2* deletion affected the Snf2 enrichment over the *ERO1* promoter. Besides *ada2Δ/Δ* and *ume6Δ/Δ*, the *set2Δ/Δ* has a negligible effect on Snf2 recruitment. Furthermore, Snf2 binding over the *FKB2* promoter was not affected in any of these mutants (Figure 2A), suggesting a histone PTM independent recruitment of SWI/SNF over the *FKB2* promoter.

**Figure 2.**
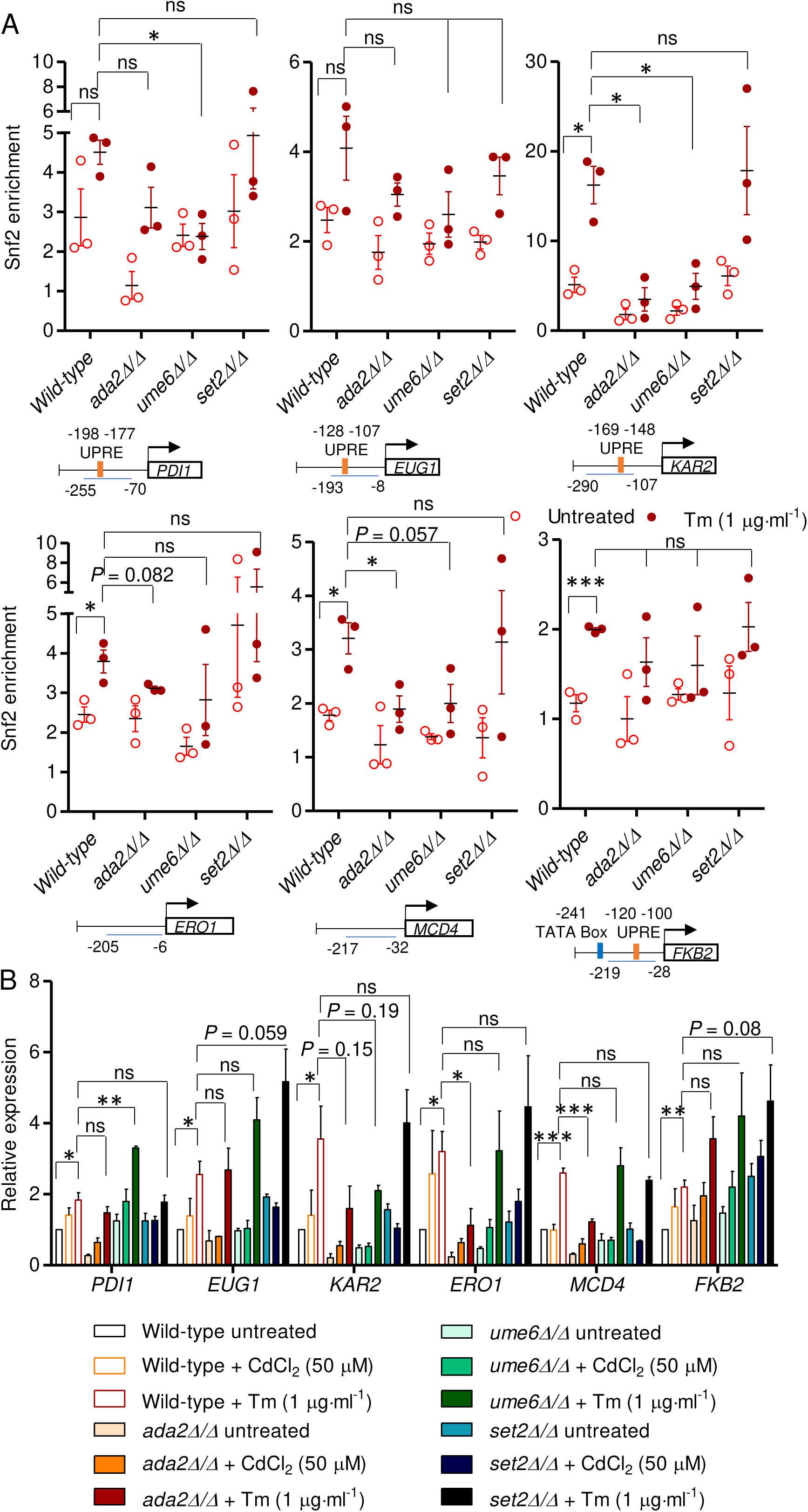
SWI/SNF binding over UPR gene promoters depends on the activity of Ada2 and Ume6. (A) The Snf2 ChIP showing fold enrichment of Snf2 over UPR gene promoters in wild-type, *ada2Δ/Δ*, *ume6Δ/Δ* and *set2Δ/Δ* cells under unstressed and tunicamycin stress conditions. The values show the Snf2 enrichment and the mean ± SEM of three independent biological repeats (n = 3). Student’s *t*-test statistical analyses were performed, and significant differences are indicated as; *, *P* ≤ 0.05; ***, *P* ≤ 0.001; ns, non-significant. (B) The relative expression of UPR genes in wild-type, *ada2Δ/Δ*, *ume6Δ/Δ* and *set2Δ/Δ* cells under ER stress are shown. Values shown are the mean and SEM of three independent biological repeats (n = 3). Student’s *t*-test statistical analyses were performed, and significant differences are indicated as; *, *P* ≤ 0.05; **, *P* ≤ 0.005; ***, *P* ≤ 0.001; ns, non-significant.

Although the Snf2 binding is reduced in *ume6Δ/Δ*, the expression of UPR genes is normal in this mutant except for *KAR2* (Figure 2B). The *ADA2* deletion, on the contrary, affects the expression of *PDI1*, *KAR2*, *ERO1* and *MCD4* in tunicamycin induced ER stress in comparison to the wild-type cells (Figure 2B). However, we cannot correlate this decrease in UPR gene inductions in *ada2Δ/Δ* and *ume6Δ/Δ* cells to the Snf2 occupancy defect seen in these mutants because the *SNF2* deletion itself shows normal UPR induction in ER stress (Figure 1C). Therefore, the disruption of *ADA2* and *UME6* must be affecting the recruitment of other chromatin remodelers like RSC or ISW1, which might be the reason for the decreased gene expression observed.

### SWI/SNF occupies the promoters of HSP, PQC genes, oxidative stress- and DNA damage-responsive genes in stress conditions

The cell’s protein quality control mechanisms, such as ubiquitin-proteasome mediated degradation and ribosome quality control (RQC), eliminate the unfolded proteins or misfolded nascent peptides in response to the ER stress (86). Under such stress, the expression of ubiquitin (Ubi4) and Rpn4 transcription factor, which stimulates activation of proteasome genes, increases. We found that SWI/SNF occupies the TATA box regions of *UBI4* and *RPN4* promoters in cadmium and tunicamycin mediated ER stress (Figure 3A). Further, its occupancy also increases at the *CDC48* promoter, encoding an AAA ATPase involved in the RQC and ERAD mechanisms (Figure 3A). In the RQC pathway, Cdc48 primarily functions in the extraction of polyubiquitinated nascent polypeptide chains from stalled ribosomes and facilitating their degradation by the proteasome.

**Figure 3.**
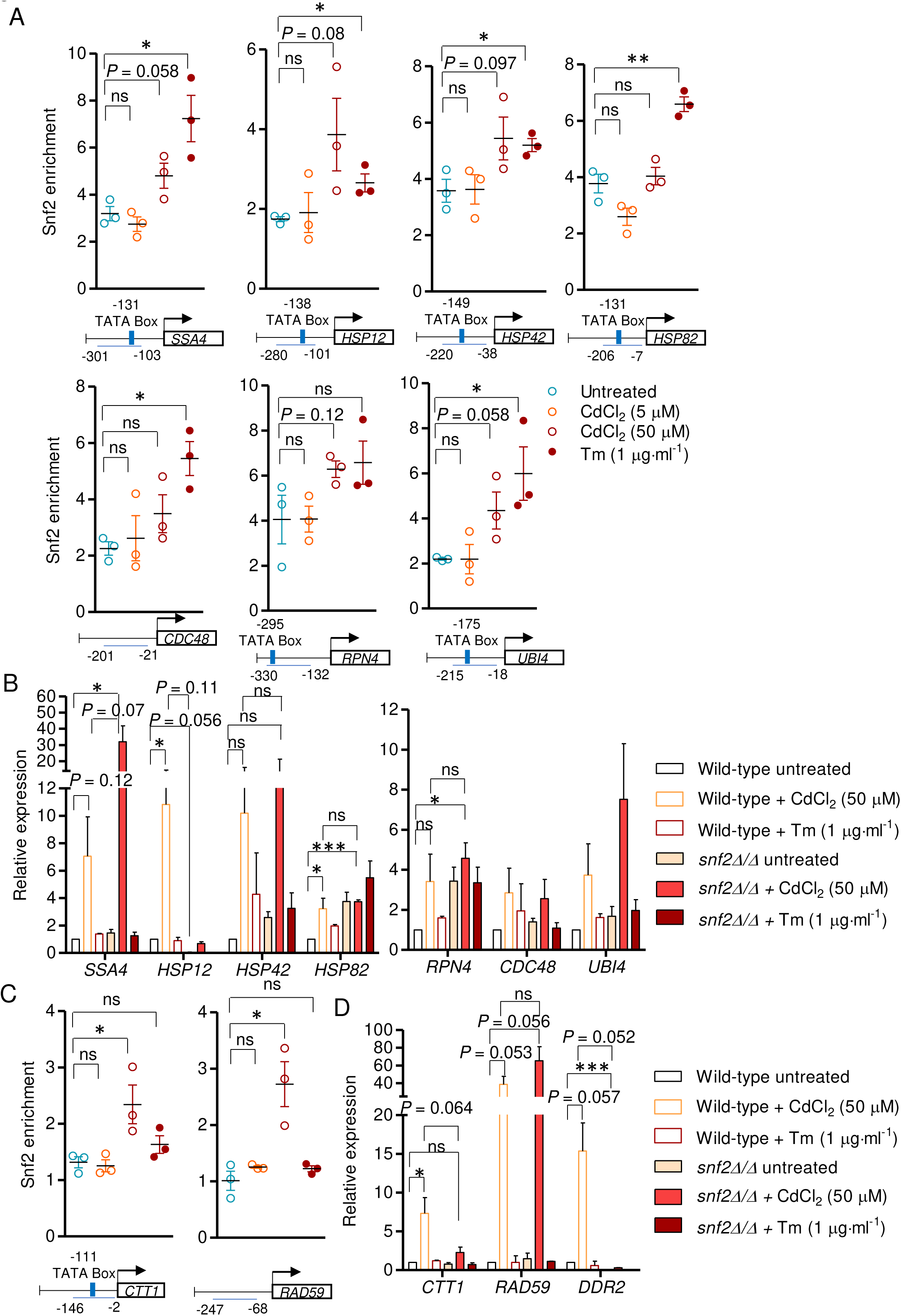
SWI/SNF occupies the promoters of HSP, PQC, oxidative stress- and DNA damage-responsive genes. (A) The Snf2 enrichment over HSP and PQC gene promoters is shown. The values indicate the Snf2 fold enrichment and the mean ± SEM of three independent biological repeats (n = 3). Student’s *t*-test statistical analyses were performed, and significant differences are indicated as; *, *P* ≤ 0.05; **, *P* ≤ 0.005; ns, non-significant. (B) The relative expression of HSP and PQC genes in wild-type and *snf2Δ/Δ* cells under ER stress are shown. Values shown are the mean and SEM of three independent biological repeats (n = 3). Student’s *t*-test statistical analyses were performed, and significant differences are indicated as; *, *P* ≤ 0.05; ***, *P* ≤ 0.001; ns, non-significant. (C) The ChIP analysis is showing the fold enrichment of Snf2 over *CTT1* and *RAD59* promoters. The values show mean enrichment ± SEM of three independent biological repeats (n = 3). Student’s *t*-test statistical analyses were performed, and significant differences are indicated as; *, *P* ≤ 0.05; ns, non- significant. (D) The relative expression of *CTT1*, *RAD59* and *DDR2* in wild-type and *snf2Δ/Δ* cells under ER stress are shown. Values shown are the mean and SEM of three independent biological repeats (n = 3). Student’s *t*-test statistical analyses were performed, and significant differences are indicated as; *, *P* ≤ 0.05; ***, *P* ≤ 0.001; ns, non-significant.

We then looked into the SWI/SNF’s involvement in some UPR independent chaperone genes and found that its enrichment increases at *SSA4* promoter in both cadmium and tunicamycin mediated ER stress (Figure 3A). While at *HSP12* and *HSP42* promoters, SWI/SNF’s enrichment increases moderately in cadmium and tunicamycin exposure (Figure 3A). The *HSP82* promoter showed an increase in binding of SWI/SNF in tunicamycin induced ER stress conditions but not in cadmium stress (Figure 3A).

These HSPs perform diverse functions inside the cell to maintain normal cellular physiology. The Hsp12 is a membrane-bound chaperone that stabilizes membranes in stress conditions. In the cytosol, Hsp12 remains unstructured, but it adopts a helical conformation in a lipid bilayer, where it influences membrane fluidity and integrity (87). The Hsp42, on the contrary, is a cytosolic chaperone, function in disaggregating the unfolded protein aggregates by forming barrel-shaped oligomers (88). Similarly, Ssa4 is a member of cytosolic HSP70 proteins, which primarily serves as a molecular chaperone for nascent polypeptides, assisting their proper folding and preventing them from aggregation (89). The *HSP82* encodes Hsp90 chaperone, which is required for folding a subset of proteins that are difficult to fold into their native conformations. Under heat stress, Hsp90 protect these proteins from thermal inactivation and enhance their refolding (90–93).

Again, to investigate the necessity of SWI/SNF for HSP and PQC genes expression, we monitored the gene induction in *SNF2* deletion strain in cadmium and tunicamycin induced ER stressed condition. We found that similar to the UPR genes, the induction of HSP and PQC genes is not affected in the *SNF2* deletion condition, except the *HSP12* (Figure 3B). Altogether these results suggest that although SWI/SNF is involved in remodelling the HSP and PQC gene promoters, its absence can be compensated by other chromatin remodelling factors. However, SWI/SNF is indispensable at *HSP12* promoter during its remodelling in cadmium stress as *SNF2* deletion drastically reduces the *HSP12* induction.

Cadmium targets multiple physiological processes, causing redox imbalance and DNA damage, evidenced by the upregulation of cytosolic catalase T (Ctt1) and proteins involved in DNA damage repair-Rad59 and Ddr2 upon cadmium treatment. We found that SWI/SNF is involved in their upregulation in cadmium stress. The SWI/SNF enrichment increases at the promoters of *CTT1* and *RAD59* post cadmium exposure (Figure 3C). In contrast, the induction of RAD59 in cadmium stress was not affected in the absence of *SNF2* (Figure 3D). The *CTT1* and *DDR2*, however, showed decreased expression in *snf2Δ/Δ* cells compared to the wild-type cells (Figure 3D). This suggests that SWI/SNF is required to remodel *CTT1* and *RAD59* promoters in cadmium-induced oxidative and DNA damaging stress, nonetheless dispensable for *RAD59* expression.

### The histone loss at UPR and other stress responsive gene promoters occur even in the absence of SWI/SNF activity

To investigate the chromatin remodelling activity at the UPR and other ER-stress responsive promoters, we performed the histone H3 ChIP. In tunicamycin treated wild-type cells, the H3 abundance at most UPR and PQC promoters gradually decreases over time, indicating nucleosome remodelling at these promoters (Figure 4A). A more significant histone displacement was detected at the *EUG1*, *KAR2*, *SSA4* and *CDC48* promoters. Similarly, the levels of H3 displacement in *snf2Δ/Δ* cells was identical to or higher than that of wild-type cells (Figure 4A). Further, in *snf2Δ/Δ* cells, at some promoters like *EUG1*, *ERO1*, *FKB2*, *SSA4* and *CDC48*, the density of H3 was much less in the unstressed condition, which might be due to a compensatory action of other chromatin remodelling factor in these cells. Overall, the H3 displacement results and the gene expression patterns are concordant.

**Figure 4.**
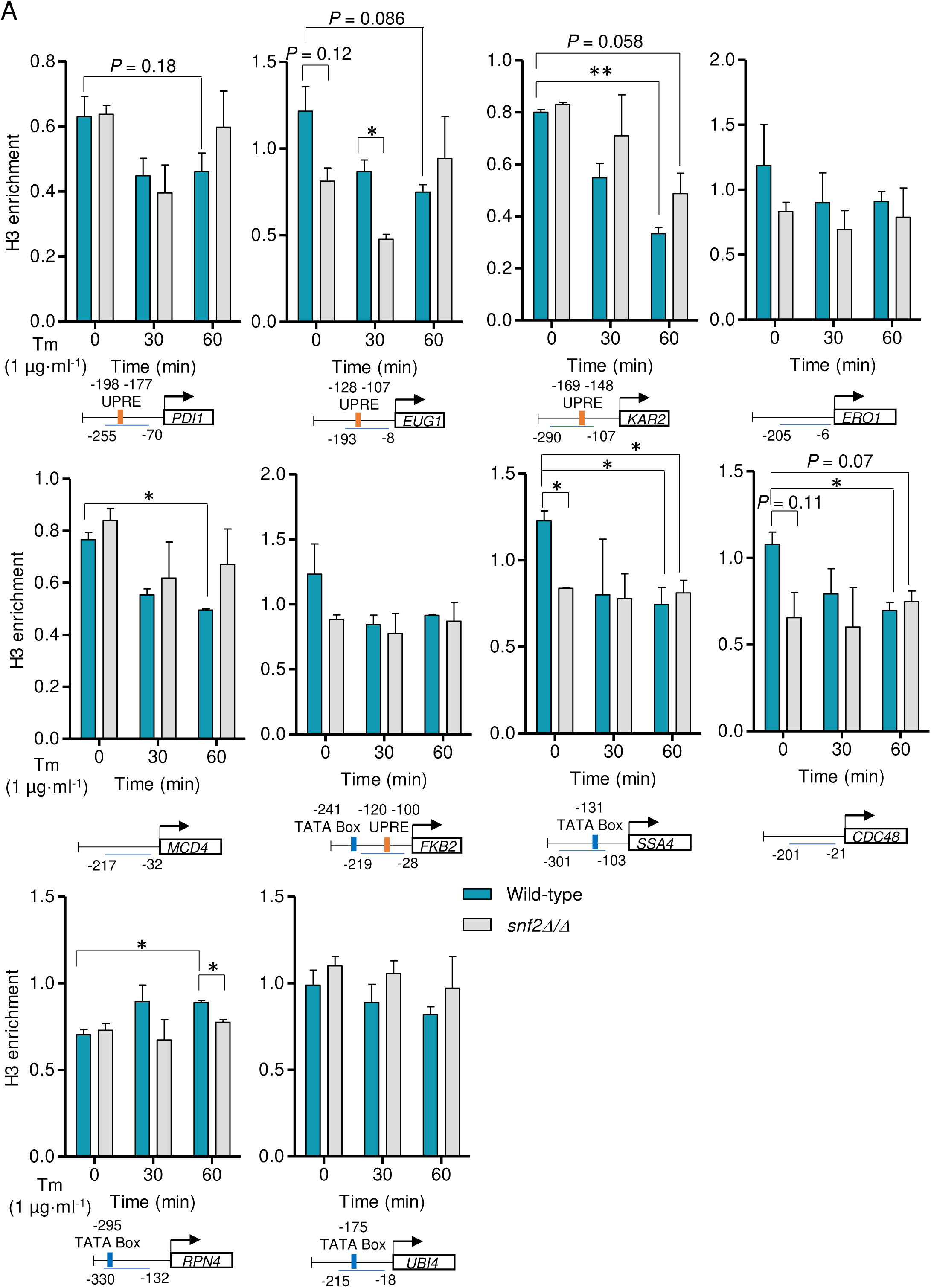
Tunicamycin induced histone loss at UPR and PQC gene promoters occurs even in the absence of SWI/SNF activity. (A) The H3 enrichment in wild-type and *snf2Δ/Δ* cells over the UPR and PQC gene promoters is shown. The values indicate the mean H3 fold enrichment and SEM of two independent biological repeats (n = 2). Student’s *t*-test statistical analyses were performed, and significant differences are indicated as; *, *P* ≤ 0.05; **, *P* ≤ 0.005; ns, non-significant.

The H3 loss was also evident in cadmium-induced ER stress at the HSP, *CDC48*, *UBI4*, *CTT1* and *RAD59* promoters (Figure 5A), which is in agreement with the gene induction pattern (Figure 5A). However, the *RPN4* promoter did not show H3 loss, but the overall H3 occupancy at this promoter was lesser than other promoters tested (Figure 4, 5), indicating that the promoter is maintained in an open chromatin conformation. This result is consistent with the high Snf2 enrichment over the *RPN4* promoter in unstressed condition (Figure 3A). Further, the decrease in H3 occupancy at these promoters was not affected in *SNF2* deletion except *HSP12*, *HSP42* and *RAD59* promoters (Figure 5A). Altogether these results suggest that although SWI/SNF is involved in chromatin remodelling at the stress-responsive promoters, its absence has a negligible effect on nucleosome displacement. Therefore, it is plausible to hypothesize that another chromatin remodelling factor acts on these promoters and compensates for the deficiency of SWI/SNF activity.

**Figure 5.**
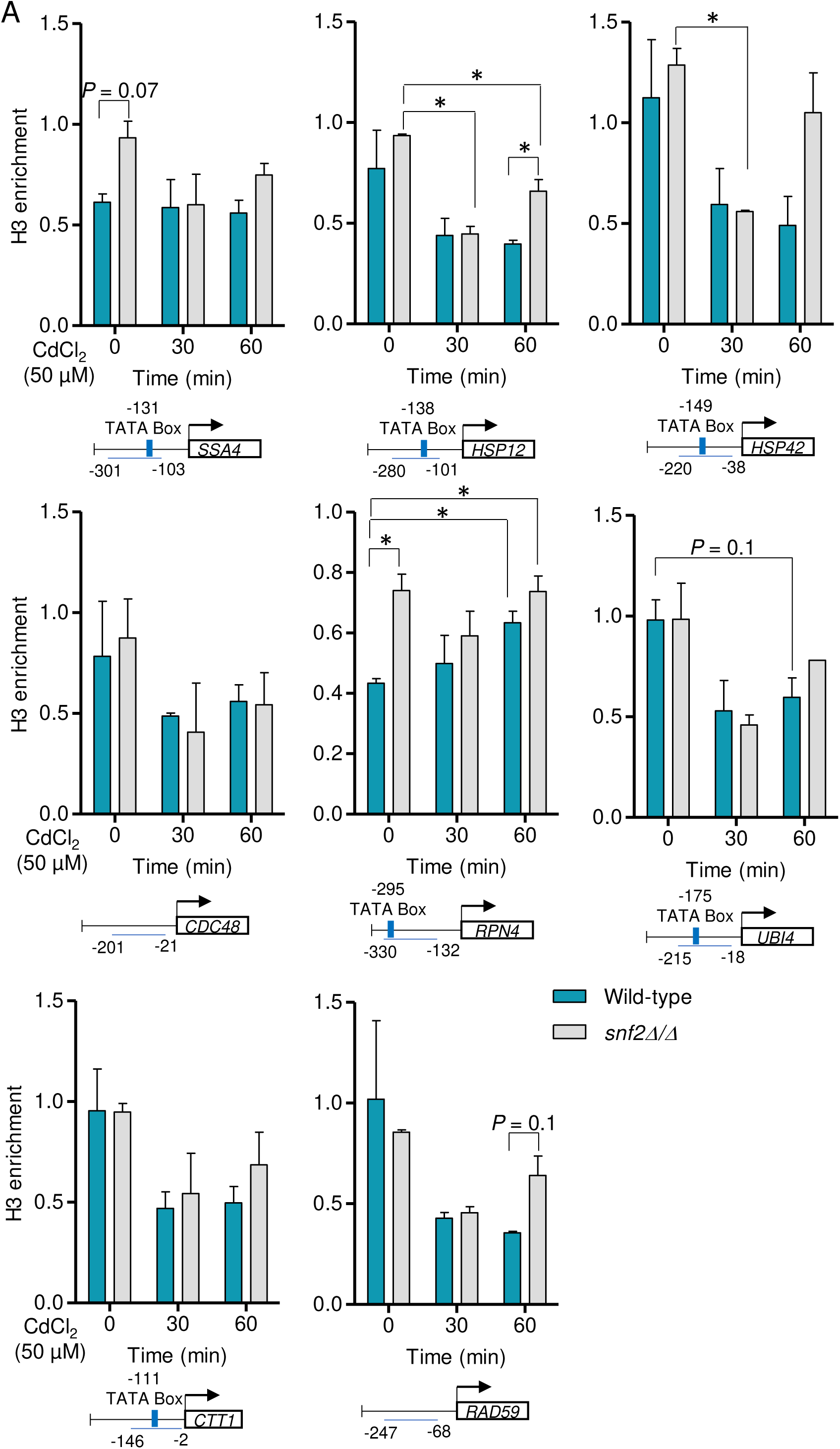
The cadmium-induced histone loss at the responsive gene promoters occurs even in the disruption of SWI/SNF activity. (A) The H3 enrichment in wild-type and *snf2Δ/Δ* cells over the promoters of HSP and cadmium responsive genes is shown. The values indicate the mean H3 fold enrichment and SEM of two independent biological repeats (n = 2). Student’s *t*-test statistical analyses were performed, and significant differences are indicated as; *, *P* ≤ 0.05; ns, non-significant.

### SWI/SNF and RSC complexes functionally cooperate to induce UPR genes in ER stress

Since the UPR gene induction in the absence of *SNF2* was similar to or higher than wild-type cells, we explored the possible involvement of the RSC complex in mediating the gene expression upon the disruption of SWI/SNF’s function. To test this, we utilized the *_TET_STH1* strains, where the promoter of *STH1* (encodes the subunit of RSC) is replaced with doxycycline (DOX) repressible promoter (63). We used this repressible system because the *STH1* knockout cells are inviable. The *STH1* expression in the *_TET_STH1* strain can be conditionally repressed by adding doxycycline in the growth media. Further, the *STH1* repression was confirmed by checking the *STH1* transcript level after 12-15 h of doxycycline supplementation in the media.

Analysis of UPR gene expression in *STH1* repression in wild-type and *snf2Δ* cells showed a functional redundancy between SWI/SNF and RSC complexes during UPR induction. The expression of *PDI1*, *EUG1* and *MCD4* in tunicamycin mediated ER stress is reduced in *snf2Δ_TET_STH1* double mutants compared to tunicamycin induced wild-type and *snf2Δ* cells (Figure 6A). Similarly, the induction of *FKB2* and *SCJ1* genes is reduced in *STH1* depleted *snf2Δ* cells compared to the wild-type, *snf2Δ* cells and in only *STH1* depletion condition (Figure 6A). Furthermore, a greater defect in gene induction was observed for *ERO1* in tunicamycin treated *snf2Δ_TET_STH1* double mutants than the *snf2Δ* and *_TET_STH1* single mutants (Figure 6A).

**Figure 6.**
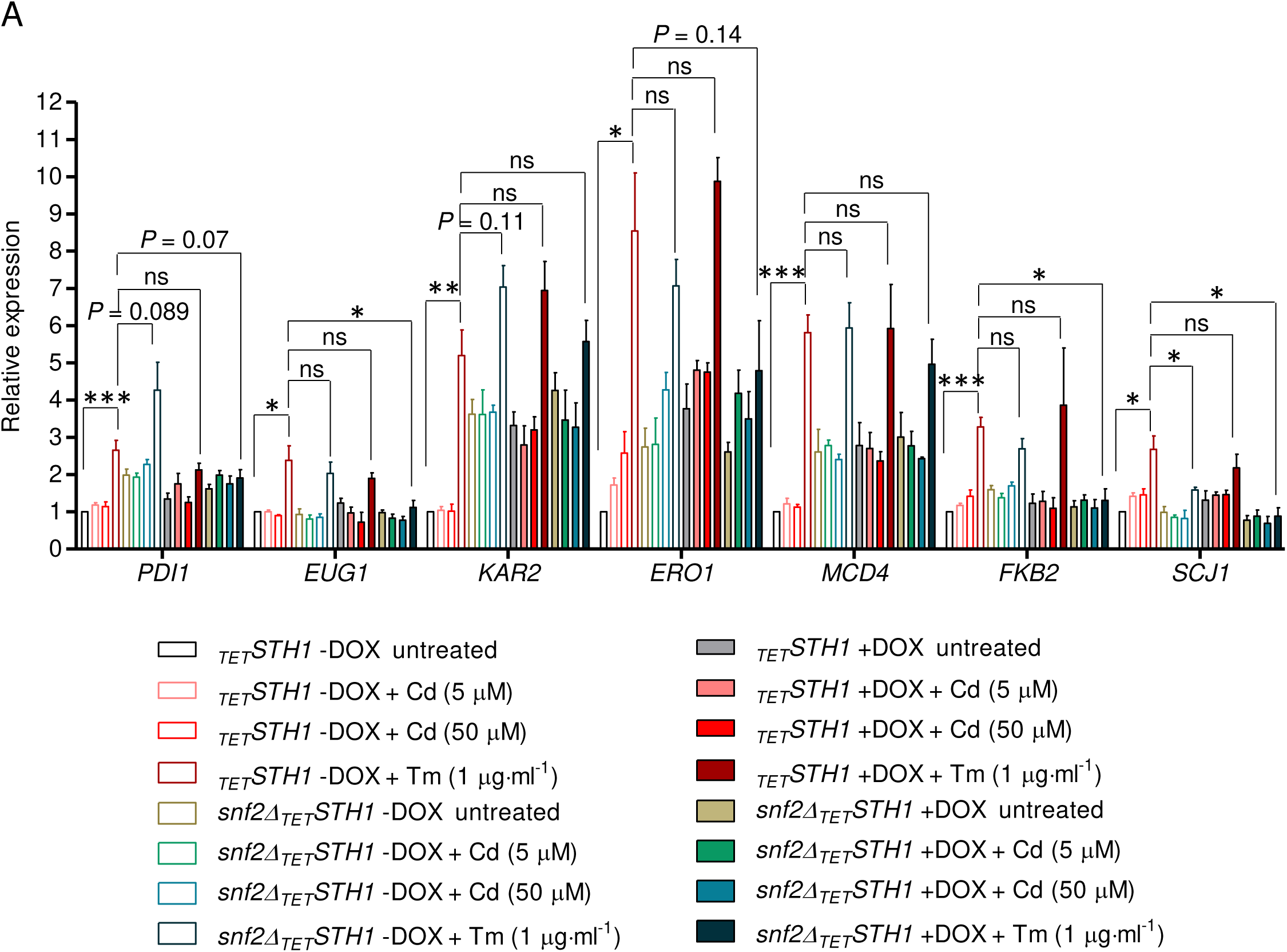
SWI/SNF and RSC complexes functionally cooperate to induce UPR genes in ER stress. (A) Expression of UPR genes in wild-type and *snf2Δ* cells without (- doxycycline) or with (+ doxycycline) *STH1* depletion. The expression of each was quantified relative to the unstressed wild-type cells without *STH1* depletion (*_TET_STH1* - DOX untreated condition). Values shown are the mean and SEM of two independent biological repeats (n = 2). Student’s *t*-test statistical analyses were performed, and significant differences are indicated as; *, *P* ≤ 0.05; **, *P* ≤ 0.005; ***, *P* ≤ 0.001; ns, non-significant.

Surprisingly, the expression of *KAR2* was not affected in *snf2Δ_TET_STH1* double mutation, suggesting a possible involvement of other remodelling factors at this promoter (Figure 6A). These results indicate the cooperative action of SWI/SNF and RSC in remodelling the UPR promoters in response to the ER stress. Additionally, a compensatory mechanism might exist between these two remodelers to ensure proper gene induction in response to the stress.

### SWI/SNF and RSC are required for the induction of HSP, *UBI4* and *RAD59* genes

We next examined the cooperative action of SWI/SNF and RSC at HSP, PQC and DNA damage responsive genes. We observed functional dependencies in SWI/SNF and RSC at *HSP12*, *HSP42* and *HSP82* but not on *SSA4* expression in response to the cadmium stress (Figure 7A). The *HSP12* expression drastically increases in the *_TET_STH1* single mutation, which may indicate that SWI/SNF compensates for the loss of Sth1 (Figure 7A). Further cooperation between SWI/SNF and RSC is also required to induce *UBI4* and *RAD59* because the expression of these genes drastically reduced in cadmium-induced *snf2Δ_TET_STH1* double mutants than wild-type cells (Figure 7B). Induction of *DDR2* is affected in *snf2Δ_TET_STH1* double mutants similar to the *snf2Δ* cells, suggesting *DDR2* requires SWI/SNF but not RSC for its induction in cadmium. On the contrary, expression of other genes such as *RPN4*, *CDC48* and *CTT1* did not indicate functional cooperation between these two remodelling complexes (Figure 7B).

**Figure 7.**
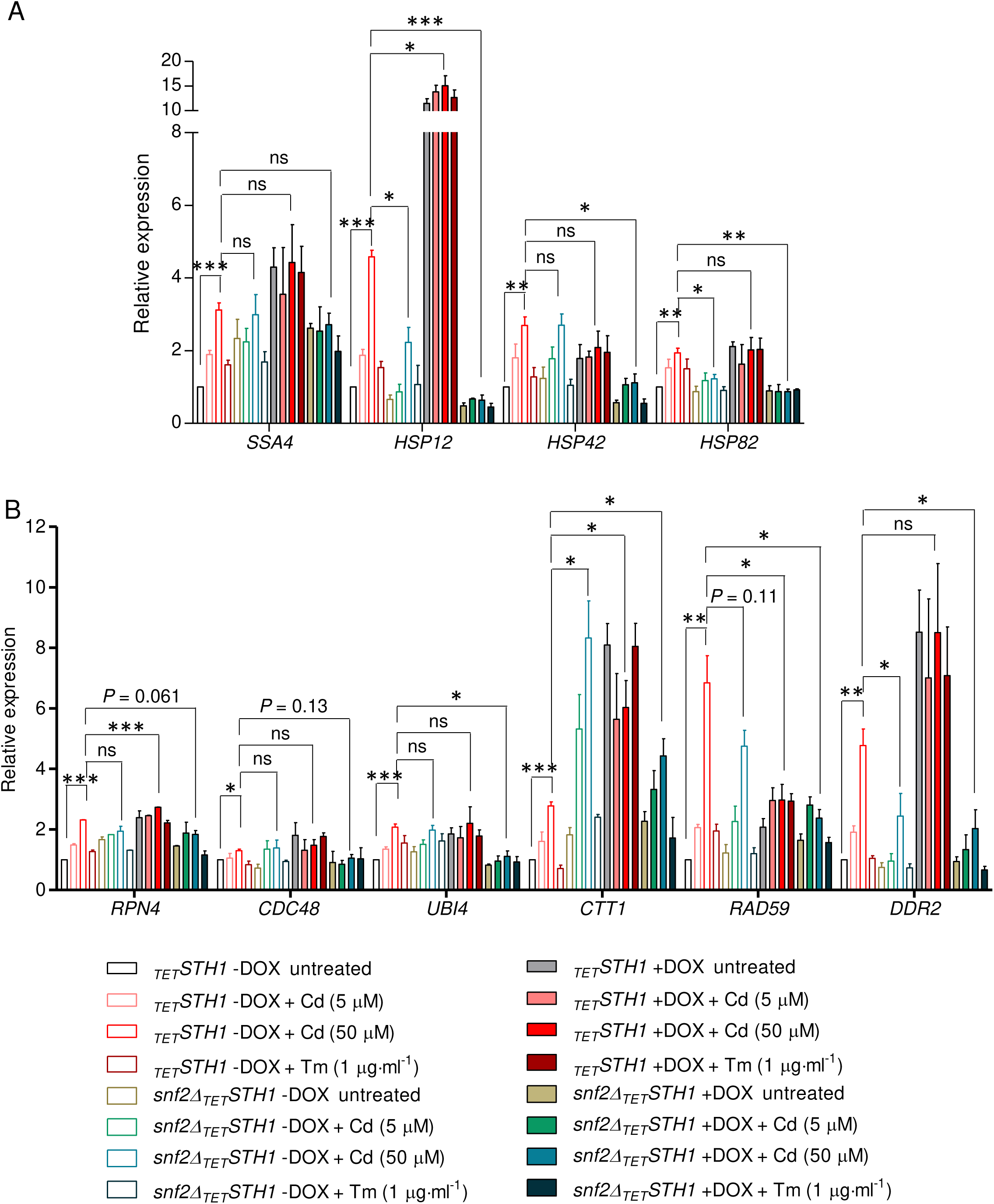
SWI/SNF and RSC are required for the induction of HSP, *UBI4* and *RAD59*. (A, B) The relative expression of HSP and other stress-responsive genes in wild-type and *snf2Δ* cells without (- doxycycline) or with (+ doxycycline) the depletion of *STH1*. Values shown are the mean and SEM of two independent biological repeats (n = 2). Student’s *t*-test statistical analyses were performed, and significant differences are indicated as; *, *P* ≤ 0.05; **, *P* ≤ 0.005; ***, *P* ≤ 0.001; ns, non-significant.

## Discussion

### A novel role of SWI/SNF in activating UPR target genes in response to the unfolded protein stress

The binding of Hac1 to its target promoters is the critical step of UPR signalling transduced by the ER-resident sensor protein Ire1. This event is accompanied by two SAGA subunits- Ada2 and Gcn5, which physically interacts with Hac1 and help to initiate the expression of its targets (94). Besides this, the involvement of other chromatin-modifying factors such as the ATP-dependent chromatin remodelling complexes in the induction of Hac1 target genes was unknown. In this study, we tested the involvement of such complexes- SWI/SNF and RSC in UPR gene induction in response to the unfolded protein stress created by tunicamycin and cadmium. Our results suggest that SWI/SNF complex occupies the promoters of UPR targets, and its enrichment increases when cells experience unfolded protein stress (Figure 1A). However, in some promoters like *PDI1*, *EUG1*, *KAR2* and *ERO1*, its binding is already high in an unstressed condition (Figure 1A, 2A), indicating SWI/SNF is involved in their basal level expression to ensure clearance of unfolded proteins generated during normal physiological processes.

Recruitment of chromatin remodelers requires a gene-specific transcriptional activator or repressor (68, 69, 85, 95). In the case of UPR genes, Hac1 acts as the SWI/SNF recruiting factor (Figure 1B); however, we still do not know the mode of its recruitment. Evidence suggests that Hac1 also interact with RPD3-SIN3 HDAC to negatively regulate the expression of early meiotic genes in nitrogen-rich growth conditions (96). Thus, Hac1 can cooperate with both activating or repressive epigenetic factors to generate different transcriptional outcomes.

The UPR gene transcription is a multifactorial event that requires activator, acetyltransferases and chromatin remodelers. The SWI/SNF binding may require additional chromatin factors. To find out those chromatin factors required for SWI/SNF binding, we tested its occupancy over UPR promoters in *ADA2*, *UME6* and *SET2* deleted strains. Snf2 binding was compromised in *ADA2*, and *UME6* deleted strain on UPR gene promoters (*KAR2*, *ERO1*, *EUG1* and *MCD4*) (Figure 2A). The reduced Snf2 binding observed in *ADA2* deleted cells was correlated with the reduced expression of *KAR2*, *ERO1*, *MCD4*, *PDI1* in Tm induced stress condition (Figure 2B). In contrast, *UME6* deletion does not reduce gene expression levels except *KAR2* (Figure 2B). Based on our results, we can speculate the chain of events at UPR gene induction, where Ada2 and Ume6 maintain a prerequisite acetylated state of the nucleosomes at the UPR promoters, favouring the binding of SWI/SNF on to these promoters (Figure 8). The Hac1 might mediate SWI/SNF recruitment by interacting with Ada2 and Gcn5 and probably with SWI/SNF.

**Figure 8.**
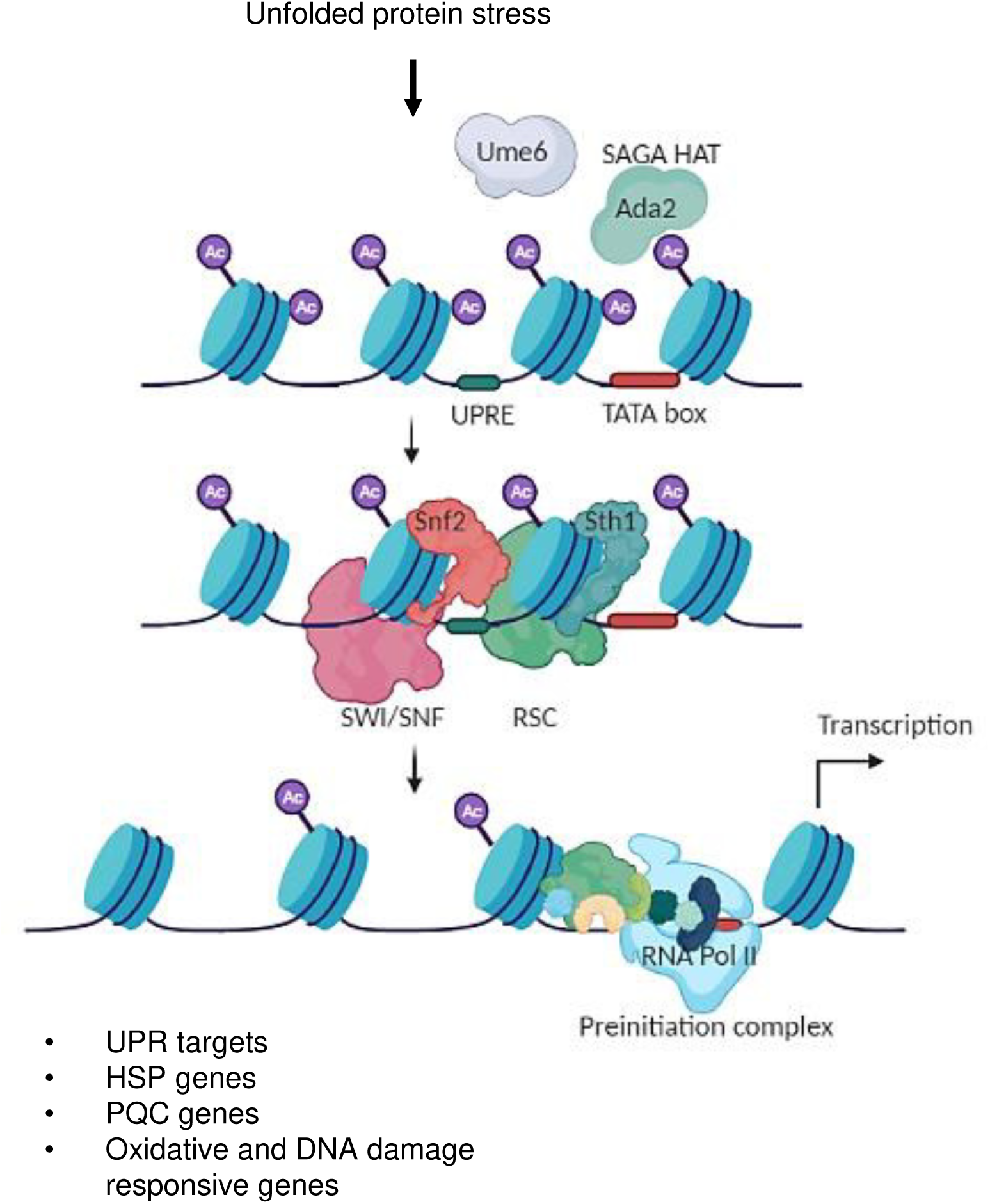
Cooperation between SWI/SNF and RSC during the induction of UPR targets and other stress responsive genes. When cell experiences unfolded protein stress, it activates genes involved in UPR, HSP, PQC, oxidative and DNA damage response pathways. Ada2 and Ume6 promote SWISNF recruitment over UPR target genes. Activation of the majority of UPR, HSP and other stress responsive genes requires the cooperative action of SWI/SNF and RSC. These complexes remodel the promoter regions to promote preinitiation complex formation and transcription initiation.

### SWI/SNF is involved in the regulation of stress-induced expression of HSP and PQC genes

Although the involvement of SWI/SNF in remodelling the promoters of *HSP12*, *HSP82* and *SSA4* in heat shock is extensively studied (64), its role in the induction of HSP and PQC genes upon cadmium and tunicamycin induced ER stress was unexplored. We have demonstrated that SWI/SNF binds to HSP and PQC gene promoters in cadmium and tunicamycin stress; however, the level of induction of these genes differs in these stresses. The tunicamycin and cadmium have a different mode of toxicity. While tunicamycin blocks the initial step of N-linked protein glycosylation, cadmium targets the nascent peptides in the ER lumen for aggregation, probably by binding to their exposed thiol groups (2, 97). Therefore, possibly, the cells accumulate a higher number of nascent peptide aggregation in cadmium than tunicamycin, demanding the activation of disaggregating chaperones and degradation machinery. This hypothesis was supported by the observation that all HSP and PQC genes were comparatively more expressed in cadmium than in tunicamycin stress.

Further cadmium targets multiple pathways in the cells, including redox homeostasis and DNA damage response. Our ChIP analysis showed the involvement of SWI/SNF in remodelling the promoters of HSP (including *SSA4* and *HSP12*), PQC genes, *CTT1* and *RAD59* in cadmium stress (Figure 3A, C). However, SWI/SNF also occupies HSP and PQC gene promoters in tunicamycin stress (Figure 3A, C). These results indicate that SWI/SNF has a stress-dependent promoter preference, although the signalling for its promoter selection is not well understood. Further biochemical assays are required to understand the binding preference of SWI/SNF in different stresses.

### SWI/SNF is not an absolute requirement for the remodelling activity at UPR, HSP and PQC gene promoters

Despite the binding of SWI/SNF at UPR, HSP and PQC genes, the *SNF2* deleted cells expressed these genes in normal levels compared to the wild-type cells (Figure 1C, 3B, D). Therefore, to confirm the chromatin remodelling activity in *SNF2* deleted cells, we tested the histone H3 depletion. We performed histone H3 ChIP to determine total H3 abundance at these promoters upon tunicamycin and cadmium-induced stress. Gene expression generally correlates with a reduction in H3 abundance or histone content. A gradual decrease in histone H3 content at UPR, HSP and PQC gene promoters suggests that histone H3 loss is evident post ER stress in wild-type cells (Figure 4, 5). Furthermore, in *SNF2* deleted cells, at the majority of these promoters the histone H3 content was similar to wild-type cells suggesting the deletion of *SNF2* does not hamper histone loss (Figure 4, 5). A similar observation was also reported in *SNF2* deleted cells at the *HSP82* promoter, suggesting that although SWI/SNF acts on the *HSP82* promoter in response to heat shock, remodelling of this promoter occurs efficiently even in its inactivation (44).

### Functional cooperation between SWI/SNF and RSC is required to express UPR and HSP genes in ER stressed condition

Different chromatin remodelers interact in a cooperative or antagonistic manner to modulate the transcriptional output. Previous studies have reported many such relationships between the different family of chromatin remodelers. One such example is the antagonism between RSC and ISWI complexes. Generally, ISWI forms regular spaced nucleosomal arrays, whereas RSC performs disorganization or ejection of nucleosomes. RSC allows gene activation by maintaining NDRs at promoters, and ISW1a counteracts this function by its repositioning activity (98). Another study suggested the antagonism between SWI/SNF and ISW2 at *RNR3* gene promoter wherein deletion of ISW2 leads to constitutive recruitment of SWI/SNF complex and overexpression of ISW2 reduces SWI/SNF binding at *RNR3* promoter (99). Similar studies have demonstrated the functional interaction by inactivating individual or multiple remodelers in combination. Elimination of *SNF2* alone slightly delayed H3 loss at *HSP82* and *SSA4* in heat shock, whereas abolishing the remodelling of *HSP12* promoter. In contrast, conditional depletion of *STH1* leads to diminished and delayed histone loss in addition to reduction in Pol II recruitment at all three promoters. Furthermore, the inactivation of *SNF2* and *ISW1* together have a synergistic effect on histone displacement (65).

Previously it has been shown that SWI/SNF and RSC work cooperatively in driving nucleosome eviction at highly expressed sulfometuron methyl induced and constitutively induced genes (63). They tested the H3 and H2B occupancies by ChIP-seq upon deleting *SNF2* and depleting Sth1. We have also tried to search for such functional interplay between SWI/SNF and RSC remodeler at UPR and other stress responsive gene promoters. To dissect the involvement of some other chromatin remodeler, we depleted *STH1* in the *SNF2* deleted background. The absence of both the ATPase subunits (Snf2 and Sth1) diminished the amplitude of gene expression compared to the wild-type and *snf2Δ* cells (Figure 6, 7). This suggests that these two remodelers probably work together (Figure 8), and the remodelling is less efficient when both are absent. Thus, both SWI/SNF or RSC are required for gene transcription at some genes, whereas in many instances (e.g., at *ERO1*, *MCD4* and *FKB2* promoters), either of them is sufficient for remodelling and gene expression.

Taken together, our study provides evidence for a functional cooperative relationship between SWI/SNF and RSC in inducing stress-responsive genes. This uncovers the function of chromatin-modifying factors in remodelling UPR gene promoters when the cell initiates the UPR signalling. In addition, our results demonstrate distinct requirements of SWI/SNF for remodelling selective genes in a stress-dependent manner. Further studies exploring the involvement of SWI/SNF and RSC on a genome-wide scale will expand our understanding of the stress-responsive transcriptional program.

## Materials and method

### Yeast strains

The knockout strains- *snf2Δ/Δ, ada2Δ/Δ, set2Δ/Δ, ume6Δ/Δ, hac1Δ/Δ* and *rpn4Δ/Δ*, are isogenic to BY4743 (*MATa/α his3Δ1/his3Δ1 leu2Δ0/leu2Δ0 LYS2/lys2Δ0 met15Δ0/MET15 ura3Δ0/ura3Δ0*). The *snf2Δ*, *_TET_STH1* and *snf2Δ_TET_STH1* strains are isogenic to BY4741 (*MATa his3Δ1 leu2Δ0 met15Δ0 ura3Δ0*), generously gifted by Alan G. Hinnebusch. All the yeast strains used in this study are listed in Table 1. Yeast cells were grown at 30°C in synthetic complete media (SC) containing 0.18% amino acids, adenine and uracil mix, 0.17% yeast nitrogen base, 0.5% ammonium sulfate and 2% glucose. Reagents used in media were purchased from Sigma, Himedia and BD.

**Table 1.** *Saccharomyces cerevisiae* strains used in this study

| Strain | Genotype | Source or reference |
| --- | --- | --- |
| Wild-type<br>(BY4743) | *MATa/α his3Δ1/his3Δ1 leu2Δ0/leu2Δ0*<br>*LYS2/lys2Δ0 met15Δ0/MET15*<br>*ura3Δ0/ura3Δ0* | Yeast knockout collection, Open Biosystems (YKO-OB) |
| *snf2Δ/Δ* | BY4743, *snf2Δ::KanMX* | YKO-OB |
| *ada2Δ/Δ* | BY4743, *ada2Δ::KanMX* | YKO-OB |
| *set2Δ/Δ* | BY4743, *set2Δ::KanMX* | YKO-OB |
| *ume6Δ/Δ* | BY4743, *ume6Δ::KanMX* | YKO-OB |
| *hac1Δ/Δ* | BY4743, *hac1Δ::KanMX* | YKO-OB |
| Wild-type<br>(BY4741) | *MATa his3Δ1 leu2Δ0 met15Δ0 ura3Δ0* | Open Biosystems |
| *snf2Δ* (F748) | BY4741, *snf2Δ::KanMX4* | (63) |
| *TETSTH1* | BY4741, *HIS3MX6::TETSTH1* | (63) |
| *snf2ΔTETSTH1* | F748, *HIS3MX6::TETSTH1* | (63) |

### cDNA preparation and real-time (RT) PCR

The secondary culture cells were harvested with or without the treatment of cadmium or tunicamycin for 1 h. For *STH1* depletion, the secondary cultures of *_TET_STH1* and *snf2Δ_TET_STH1* cells (at ∼0.2 OD_600_) were treated with 10 µg·ml^-1^ doxycycline for 12-15 h before the cadmium or tunicamycin insult. The total RNA isolated from the untreated or treated cells by the hot-phenol method (100) and ∼1 µg of RNA was subjected to reverse transcription PCR using iScript cDNA synthesis kit (Bio-Rad cat. No. 1708891).

We used the KAPA SYBR FAST qPCR-Kit (KAPA, cat. No. KK4618), TB Green Premix Ex Taq™ II (Takara, cat. No. RR820B) and followed the manufacturer’s standard protocol to perform RT PCR. The RT-PCR reaction was analyzed in the ABI-7300 RT-PCR machine (Applied Biosystems, CA, USA). The level gene transcripts were normalized with the *ACT1* transcript level, and a relative fold change in gene expression was calculated as the 2*^-ΔΔCT^* method (101).

### Chromatin immunoprecipitation

ChIP was performed as previously described (102), with slight modifications. Cells were grown up to mid-log phase (∼0.7-0.8 OD_600_) and treated with cadmium or tunicamycin for 1 h, followed by cross-linking with 1% formaldehyde for 12-15 min at 25°C and quenching with glycine for 5 min. Then cells were harvested, washed, and resuspended in FA lysis buffer (50 mM HEPES, 150 mM NaCl, 2 mM EDTA, 1% Triton X-100, 0.1% sodium deoxycholate, 0.1% SDS, protease inhibitor cocktail [PIC], and phenylmethylsulphonyl fluoride [PMSF]) and mechanically lysed by vigorous vertexing with 0.5 mm glass beads for 10 cycles of 30 sec on/off (30 sec vertexing and 30 sec incubation in ice). Cells were again vortexed for 30 min at 4°C. Chromatin sonication was carried out using a Diagenode Bioruptor, and the average size of sonicated fragments was ∼500 bp for SWI/SNF ChIP and ∼300 bp for histone H3 ChIP. After sonication, the lysate was cleared by centrifuging at 13000 rpm at 4°C for 30 min. Chromatin immunoprecipitation was carried out overnight at 4°C by incubating chromatin with 30 µl of Dynabead protein G slurry (Dynabead protein G, cat. no. 10003D; Thermo Fisher Scientific) conjugated with anti-Snf2 (gifted from Joseph C. Reese) or anti-histone H3 (Sigma-Aldrich, H0164) antibody. An equal volume of chromatin was also taken as input control.

The immunoprecipitated chromatin (IP) were washed under stringent conditions using the following buffers: low-salt buffer (0.1% Triton X-100, 2 mM EDTA, 0.1% SDS, 150 mM NaCl, 20 mM HEPES), high-salt buffer (0.1% Triton X-100, 2 mM EDTA, 0.1% SDS, 500 mM NaCl, 20 mM HEPES), LiCl buffer (0.5 M LiCl, 1% NP-40, 1%, Sodium deoxycholate, 100 mM Tris-Cl [pH 7.5]), and 1X Tris-EDTA (TE) buffer (pH 8). Then de-cross-linking was performed at 65°C overnight. DNA isolation was carried out from both IP and input samples as described previously (103).

The IP and input DNA were quantified in RTPCR using SYBR green mix (KAPA Biosystems; cat. no. KK4618). The Snf2 enrichment was normalized to the amount of input DNA and the background noise (signal derived from chromosome V intergenic region [IGR]). The enrichment was calculated using the formula; fold enrichment = 2^−*ΔΔCT*^; *ΔΔC_T_* = (*C_T promoter_* [IP] – *C_T promoter_* [input]) – (*C_T ChV IGR_* [IP] – *C_T ChV IGR_* [input]), where *C_T_* is cycle threshold. The histone H3 enrichment was normalized to the input DNA and the signal derived from *PHO5* promoter using the formula; H3 fold enrichment = 2^−*ΔΔCT*^; *ΔΔC_T_* = (*C_T promoter_* [IP] – *C_T promoter_* [input]) – (*C_T PHO5 promoter_* [IP] – *C_T PHO5 promoter_* [input]). All the ChIP experiments were conducted at least twice.

## Acknowledgements

We thank Alan G. Hinnebusch for gifting us with the yeast strains. We thank Joseph C. Reese for gifting us the anti-Snf2 antibody. This work was supported by funds from the Council of Scientific and Industrial Research, Government of India (CSIR Grant No. 38(1468)/18/EMR-II) to RST and intramural funds from IISER Bhopal. RKS and SS acknowledge IISER Bhopal and CSIR, respectively, for providing fellowship support.

## Conflicts of interest

We declare no conflicts of interest.

## Author Contributions

RKS, SS, and RST conceptualized the study. RKS and SS designed, performed all the experiments and analyzed the results with RST. RKS and SS prepared all the figures, wrote the original draft and edited the manuscript with RST. All authors reviewed the results and approved the final version of the manuscript.

